# Multi-ethnic genome-wide association study of decomposed cardioelectric phenotypes illustrates strategies to identify and characterize evidence of shared genetic effects for complex traits

**DOI:** 10.1101/654012

**Authors:** Antoine R. Baldassari, Colleen M. Sitlani, Heather M. Highland, Dan E. Arking, Steve Buyske, Dawood Darbar, Rahul Gondalia, Misa Graff, Xiuqing Guo, Susan R. Heckbert, Lucia A. Hindorff, Chani J. Hodonsky, Yii-Der Ida Chen, Robert C. Kaplan, Ulrike Peters, Wendy Post, Alex P. Reiner, Jerome I. Rotter, Ralph V. Shohet, Amanda A. Seyerle, Nona Sotoodehnia, Ran Tao, Kent D. Taylor, Genevieve L Wojcik, Jie Yao, Eimear E. Kenny, Henry J. Lin, Elsayed Z. Soliman, Eric A. Whitsel, Kari E. North, Charles Kooperberg, Christy L. Avery

**Affiliations:** Gillings School of Global Public Health, University of North Carolina at Chapel Hill, Chapel Hill NC; Cardiovascular Health Research Unit, Department of Medicine, University of Washington, Seattle, WA; McKusick-Nathans Institute of Genetic Medicine, Johns Hopkins University School of Medicine, Baltimore, MD; Department of Statistics & Biostatistics, Rutgers University, New Brunswick, NJ; Department of Medicine, University of Illinois at Chicago, Chicago, IL; Institute for Translational Genomics and Population Sciences and Department of Pediatrics, Los Angeles Biomedical Research Institute at Harbor-UCLA Medical Center, Torrance, CA; Department of Pediatrics, David Geffen School of Medicine at UCLA, Los Angeles, CA; Cardiovascular Health Research Unit, Division of Cardiology, Department of Medicine, University of Washington, Seattle, WA; Division of Genomic Medicine, National Human Genome Research Institute, National Institutes of Health, Bethesda, MD; Albert Einstein College of Medicine, Bronx, NY; Fred Hutchinson Cancer Research Center, Public Health Sciences Division, Seattle, WA; Departments of Medicine and Epidemiology Johns Hopkins University, Baltimore, MD; Center for Cardiovascular Research, John A. Burns School of Medicine, Honolulu, HI; UNC Eshelman School of Pharmacy, University of North Carolina at Chapel Hill, Chapel Hill, NC; Department of Biostatistics, Vanderbilt University, Nashville, TN; Department of Biomedical Data Science, Stanford University School of Medicine, Stanford, CA; Center for Genomic Health, Icahn School of Medicine at Mount Sinai, New York City, NY; Charles Bronfman Institute of Personalized Medicine, Icahn School of Medicine at Mount Sinai, New York City, NY; Department of Genetics and Genomic Sciences, Icahn School of Medicine at Mount Sinai, New York City, NY; Department of Medicine, Icahn School of Medicine at Mount Sinai, New York City, NY; Epidemiological Cardiology Research Center (EPICARE), Wake Forest School of Medicine, Winston-Salem, NC; Carolina Center for Genome Sciences, University of North Carolina at Chapel Hill, Chapel Hill NC

## Abstract

**Background:** Published genome-wide association studies (GWAS) are mainly European-centric, examine a narrow view of phenotypic variation, and infrequently interrogate genetic effects shared across traits. We therefore examined the extent to which a multi-ethnic, combined trait GWAS of phenotypes that map to well-defined biology can enable detection and characterization of complex trait loci.

**Methods:** With 1000 Genomes Phase 3 imputed data in 34,668 participants (15% African American; 3% Chinese American; 51% European American; 30% Hispanic/Latino), we performed covariate-adjusted univariate GWAS of six contiguous electrocardiogram (ECG) traits that decomposed an average heartbeat and two commonly reported composite ECG traits that summed contiguous traits. Combined phenotype testing was performed using the adaptive sum of powered scores test (aSPU).

**Results:** We identified six novel and 87 known ECG trait loci (aSPU p-value < 5E-9). Lead SNP rs3211938 at novel locus CD36 was common in African Americans (minor allele frequency=10%) and near-monomorphic in European Americans, with effect sizes for the composite trait, QT interval, among the largest reported. Only one novel locus was detected for the composite traits, due to opposite directions of effects across contiguous traits that summed to near-zero. Combined phenotype testing did not detect novel loci unapparent by univariate testing. However, this approach aided locus characterization, particularly when loci harbored multiple independent signals that differed by trait.

**Conclusions:** Despite including one-third as few participants as the largest published GWAS of ECG traits, our study identifies multiple novel ECG genetic loci, emphasizing the importance of ancestral diversity and phenotype measurement in this era of ever-growing GWAS.

**AUTHOR SUMMARY:** We leveraged a multiethnic cohort with precise measures of cardioelectric function to identify novel genetic loci affecting this complex, multifaceted phenotype. The success of our approach stresses the importance of phenotypic precision and participant diversity for future locus discovery and characterization efforts, and cautions against compromises made in genome-wide association studies to pursue ever-growing sample sizes.

## INTRODUCTION

Genetic susceptibility underlies a majority of common diseases and traits, shown by genome-wide association studies (GWAS) that have identified thousands of genetic loci for traits such as cardiovascular, cardiometabolic, cancer, kidney, psychiatric, ocular, inflammatory, and neuromuscular conditions(1). The many GWAS reports have revealed both common threads underlying the genetic architecture of complex diseases and traits, as well as research gaps. For example, evidence of shared genetic effects (i.e., pleiotropy) is widespread, even for traits with few known etiologic links(2,3). Yet few studies have systematically examined evidence of shared genetic effects, thereby missing opportunities to identify and characterize master regulators as strong candidates for intervention(2,4). There is also limited racial/ethnic diversity, inasmuch as the majority (>80%) of GWAS participants have been of European ancestry^3^. Limited diversity leads to a biased view of human variation that hinders translation of genetic associations into clinical and public health applications for all populations(5,6). Further, the scale and collaborative nature of current GWAS analyses prioritize traits that are widely available across studies, which may not precisely capture phenotypic variation and its underlying biology(7,8). Together, these research gaps argue for expanding GWAS analyses to systematically assess for shared genetic effects across a spectrum of biologically important traits in multi-ethnic populations.

Electrocardiograms (ECG) measure a sequence of distinct electrophysiologic processes in the myocardium that underlie cardiac conduction and repolarization. ECG traits have high heritability(9), are relevant to cardiovascular health,(10) and allow opportunities for dense phenotyping(11). Moreover, there are few racially/ethnically diverse GWAS reports on ECG traits(12). Therefore, ECG traits are well suited for assessing the degree to which increased racial/ethnic diversity, evaluation of genetic effects shared across phenotypes, and improved phenotype resolution can enhance locus identification and characterization. We therefore examined individual and shared genetic effects underlying six contiguous measures of the ECG waveform spanning an average heartbeat. Analyses used data from the multi-ethnic Population Architecture using Genetic Epidemiology (PAGE) study and the Multi-Ethnic Study of Atherosclerosis (MESA). Our results illustrate the broad utility of multi-ethnic GWAS analyses of carefully constructed individual and aggregate traits to illuminate the biology of complex diseases and traits.

## RESULTS

### Sample description

Of the 39,538 participants with GWAS and ECG data in ARIC, HCHS/SOL, MESA, and WHI, 34,668 (88%) met all inclusion criteria (Tables S1, S2). Seventy-five percent of eligible participants were female, the mean age was 55 years, and nearly half were either Hispanic/Latino (30%) or African American (15%) (Table S4). On average, participants were overweight (BMI mean = 29 kg/m^2^) and had high serum low-density lipoprotein cholesterol (mean = 135 mg/dL). There was a high prevalence of hypertension (49%). Holding all adjustment variables constant, P wave and TP segment durations were the most strongly correlated among the six ECG traits (partial correlation ρ=−0.65), whereas T wave and P wave durations (ρ=0.01) as well as QRS and PR durations (ρ=−0.02) were largely uncorrelated (Table S5).

### Overview of association results

Approximately 22M SNPs met our inclusion criteria and were used to assess for associations with the six contiguous and two composite ECG traits (Figures S1, S2). Lead SNPs at 82 of 149 loci (56%) previously reported by 26 interval scale ECG trait GWAS analyses (Table S6) were identified at genome-wide significance in our multi-ethnic population. The successful identification of previously recognized loci varied by trait (Table S7), ranging from 21 of 45 (47%) loci for QRS interval, to nine of 14 (64%) loci for P wave. When using a lower significance threshold of p_aSPU_<0.0003 (0.05/149), 123 of the 149 (83%) previously recognized interval scale ECG trait loci were identified.

An additional six loci were >500 kb away from all lead SNPs previously reported by interval scale ECG trait GWAS and are presented as novel (Table 1, Figure 1). As described below, our results highlight the utility of phenotype decomposition, ancestral diversity, and combined-phenotype testing for the identification and characterization of complex trait loci.

**Table 1.** Novel genome wide-significant (p_aSPU_ < 5×10^−9^) loci discovered in genome-wide association study of six contiguous electrocardiographic traits that decompose an average heartbeat in N=34,668 participants from the multi-ethnic Population Architectures using Genomics and Epidemiology study and the Multi-Ethnic Study of Atherosclerosis

| Lead SNP | Cnr | Position | Coded allele | Non-coded allele | Locus | AA | EA | CHN | HIS | Contiguous ECG traits |  |  |  | Composite ECG traits |  |  |  |
| --- | --- | --- | --- | --- | --- | --- | --- | --- | --- | --- | --- | --- | --- | --- | --- | --- | --- |
|  |  |  |  |  |  |  |  |  |  | P wave | PR segment | QRS interval | ST segment | T wave | TP segment | QT interval | PR interval |
| rs13143308 | 4 | 111714419 | T | G | *PITX2* | 30% | 21% | 74% | 39% | **2x10<sup>-11</sup>** | 5x10<sup>-4</sup> | 2x10<sup>-1</sup> | 3x10<sup>-2</sup> | 3x10<sup>-1</sup> | 1x10<sup>-1</sup> | 4x10<sup>-1</sup> | 8x10<sup>-1</sup> |
| rs4340921 | 5 | 49687697 | C | T | *EMB* | 66% | 46% | 49% | 44% | **8x10<sup>-13</sup>** | 8x10<sup>-1</sup> | 1x10<sup>-3</sup> | 7x10<sup>-1</sup> | 2x10<sup>-1</sup> | 2x10<sup>-4</sup> | 7x10<sup>-1</sup> | 2x10<sup>-3</sup> |
| rs3211938 | 7 | 80300449 | G | T | *CD36* | 10% | <0.01% | <0.01% | 1% | 2x10<sup>-5</sup> | 8x10<sup>-3</sup> | 5x10<sup>-3</sup> | 4x10<sup>-1</sup> | 1x10<sup>-5</sup> | **1x10<sup>-13</sup>** | **6x10<sup>-10</sup>** | 6x10<sup>-6</sup> |
| rs11073663 | 15 | 85260268 | A | G | *ZNF592* | 27% | 54% | 19% | 48% | 4x10<sup>-1</sup> | **3x10<sup>-10</sup>** | 5x10<sup>-1</sup> | 4x10<sup>-1</sup> | 2x10<sup>-3</sup> | 7x10<sup>-3</sup> | 2x10<sup>-2</sup> | 6x10<sup>-7</sup> |
| rs142166837 | 17 | 57471022 | C | T | *YPEL2* | 31% | 52% | 32% | 49% | 5x10<sup>-2</sup> | 6x10<sup>-1</sup> | 1x10<sup>-3</sup> | 1x10<sup>-1</sup> | **4x10<sup>-11</sup>** | 1x10<sup>-2</sup> | 4x10<sup>-7</sup> | 8x10<sup>-1</sup> |
| rs13047360 | 21 | 28851580 | G | A | *BC043580* | 7% | 17% | 23% | 16% | 7x10<sup>-1</sup> | 3x10<sup>-2</sup> | **2x10<sup>-11</sup>** | 2x10<sup>-1</sup> | 5x10<sup>-2</sup> | 1x10<sup>0</sup> | 2x10<sup>-1</sup> | 3x10<sup>-2</sup> |

aSPU: adaptive sum of powered tests AA: African American, CHN: Chinese-American, EA: European-American, HIS: Hispanic/Latinos Bolded values exceed the genome-wide significance threshold ( $p < 5 \times 10^{-9}$ )

**Figure 1.**
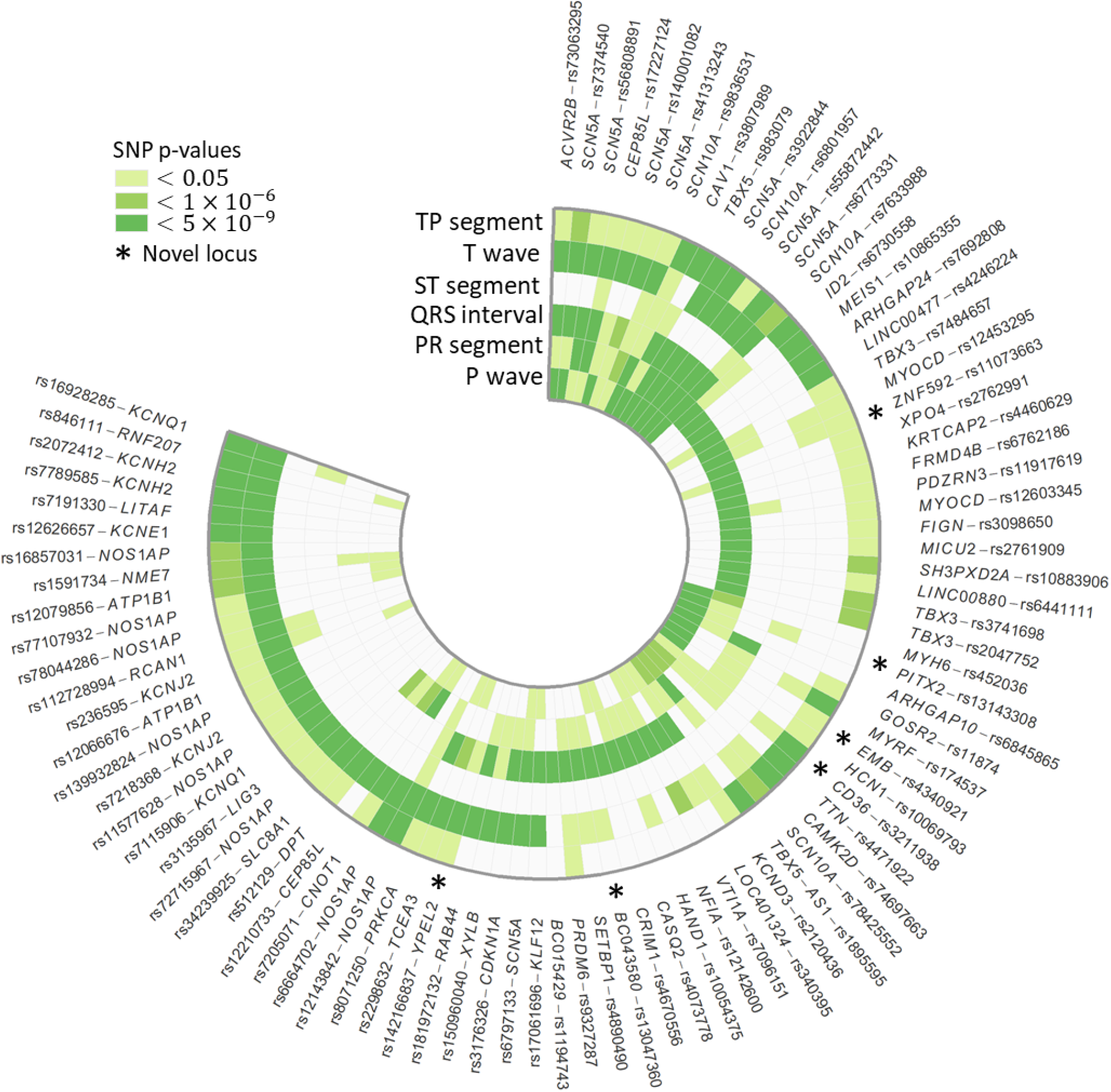
Lead SNPs at 87 loci significantly associated (p_aspu_<5×10^−9^) with six contiguous ECG traits spanning an average heartbeat, in n=34,668 multi-ethnic participants in the Population Architecture Using Genomics and Epidemiology study and Multi-Ethnic Study of Atherosclerosis. Outer stars denote novel loci and darker shades of green indicate lower p-values. To aid interpretation, lead SNPs were organized into broadly similar groups using hierarchical cluster analysis.

### Phenotype decomposition

Of the six novel loci identified for the contiguous traits, two loci were identified for P wave and one locus each was identified for PR segment, QRS interval, T wave, and TP segment. None of the novel loci were associated with ST segment or affected multiple contiguous ECG traits at genome-wide significance levels.

We then contrasted results for the six contiguous ECG traits with results from the two composite ECG traits, QT interval (QRS interval + ST segment + T wave) and PR interval (P wave + PR segment) (Figure 2). *CD36* was the only novel locus identified for both a contiguous (TP segment) and a composite (QT interval) ECG trait (Table 1). We also examined evidence of consistency of SNP effects by grouping traits according to whether they affected atrial (PR interval, PR segment, and P wave) or ventricular (QT interval, QRS interval, T wave, and ST segment) conduction. For atrial traits, novel loci identified for the contiguous traits had varying directions of effects (Figure 2a, Table S9), which when combined resulted in near-zero estimated effects for the composite trait. For example, every copy of the T allele for *PITX2* lead SNP rs13143308 increased P wave duration by 0.63 milliseconds [ms] (p_univariate_=2×10^−11^), but shortened the PR segment by 0.58 ms (p_univariate_=6×10^−4^). However, when evaluated together as the composite trait PR interval, every copy of the rs13143308 T allele prolonged the PR interval by 0.03 ms (p_univariate_=0.84). Similarly, among the 59 loci associated with ventricular conduction, two of the three novel loci (rs142166837 and rs13047360) had opposite effects on QRS interval and T wave duration, which did not reach genome-wide significance when summed for the composite trait QT interval (Figure 2, Table S9). There were no instances of either PR or QT interval identifying a novel locus not associated with any of the six contiguous traits at the genome-wide level.

**Figure 2.**
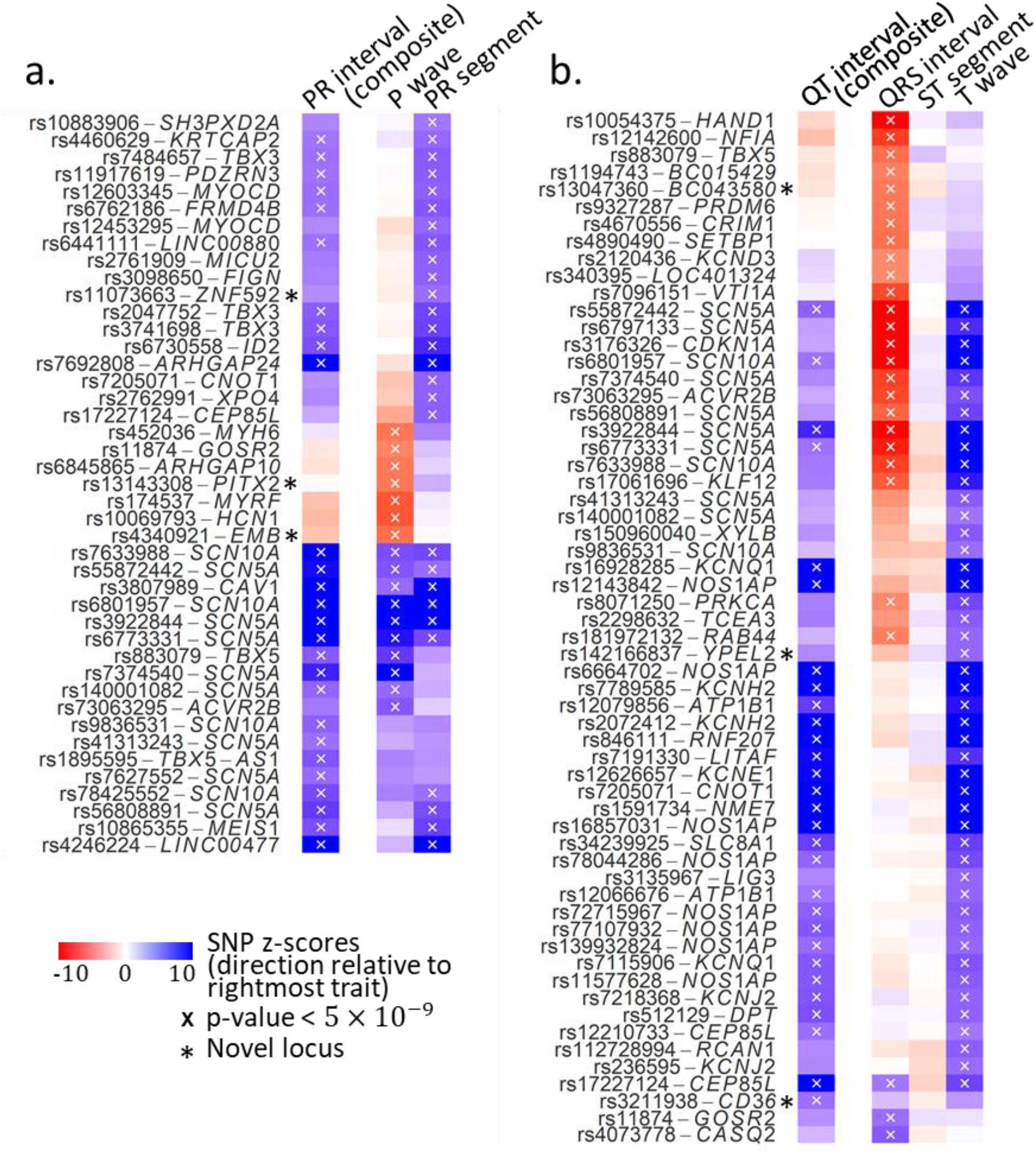
Lead SNPs at all loci significantly (P_aSPU_<5×10^−9^) associated with (a) atrial and (b) ventricular conduction measures, in n=34,668 multi-ethnic Population Architecture Using Genomics in Epidemiology (PAGE) Study and Multi-Ethnic Study of Atherosclerosis participants. Outer stars denote novel loci; blue and red shading signifies opposite directions of effect, relative to PR interval and QT interval. To aid interpretation, lead SNPs were organized into broadly similar groups using hierarchical cluster analysis.

### Ancestral diversity

Lead SNPs at five of the six novel loci were common (MAF >5%) across ancestral populations, with little evidence of heterogeneity of effect across race/ethnicity (Table S8). One locus (*CD36*) showed evidence of population specificity, with lead SNP rs3211938 near-monomorphic in European and Chinese populations (MAF<0.01%), infrequent in Hispanic/Latinos (MAF=1%), and common in African Americans (MAF=10%). Variant rs3211938 showed genome-wide significant associations with TP segment (p_univariate_=1×10^−13^) and QT interval (p_univariate_=6×10^−10^) and nominal associations with P wave (p_univariate_=2×10^−5^), PR segment (p_univariate_=0.008), and QRS interval (p_univariate_=0.005). Although no GWAS of TP segment has been published, each copy of the rs3211938 G allele increased QT interval by 3.70 ms. Reported effects for common (MAF>5%) QT lead SNPs range from 0.5 ms to 3.5 ms(13). SNP rs3211938 was either genotyped or well-imputed across studies and ancestry groups (imputation quality > 0.89, Table S10).

### Combined phenotype analyses

We found widespread evidence of shared genetic effects across ECG traits. One fourth of lead SNPs identified as genome-wide significant (P_aspu_<h5×10^−9^) had univariate associations with at least two ECG traits (p_Univariate_<5×10^−9^). Lead SNPs at *ACVR2B, SCN5A, SCN10A, CEP85L, CAV1*, and *TBX5* were associated with three or more ECG traits at univariate genome-wide significance levels. As expected, traits that were more highly correlated also showed stronger evidence of shared genetic effects, with 10 of the 20 lead SNPs that were associated with PR segment also showing genome-wide associations with TP segment. However, evidence of shared genetic effects among uncorrelated traits was also observed. For example, eight genome-wide significant SNPs at *SCN5A*, *SCN10A*, *CEP85L*, and *CNOT1* exhibited significant univariate associations with both the T wave and P wave, despite low correlation between the two traits (ρ = 0.02).

There also was evidence of allelic heterogeneity for multiple ECG traits. As an example, five signals in low LD (r^2^ < 0.1) were detected within a 500 kb region near the previously identified locus *TBX5*, each associated with a distinct combination of ECG traits. Two of the five independent signals (rs3741698 and rs2047752) were largely specific to PR segment (Figure 3). The other three signals involved PR segment and QRS interval (lead SNP rs4784657), P wave, QRS interval, and TP segment (lead SNP rs883079), and the combined phenotype only (lead SNP rs1895595). Lead SNPs also showed some evidence of variation across traits, including the locus identified by lead SNP rs7484657, for which p-values for the QRS interval lead SNP differed by approximately three orders of magnitude from the rs7484657 p-value.

**Figure 3.**
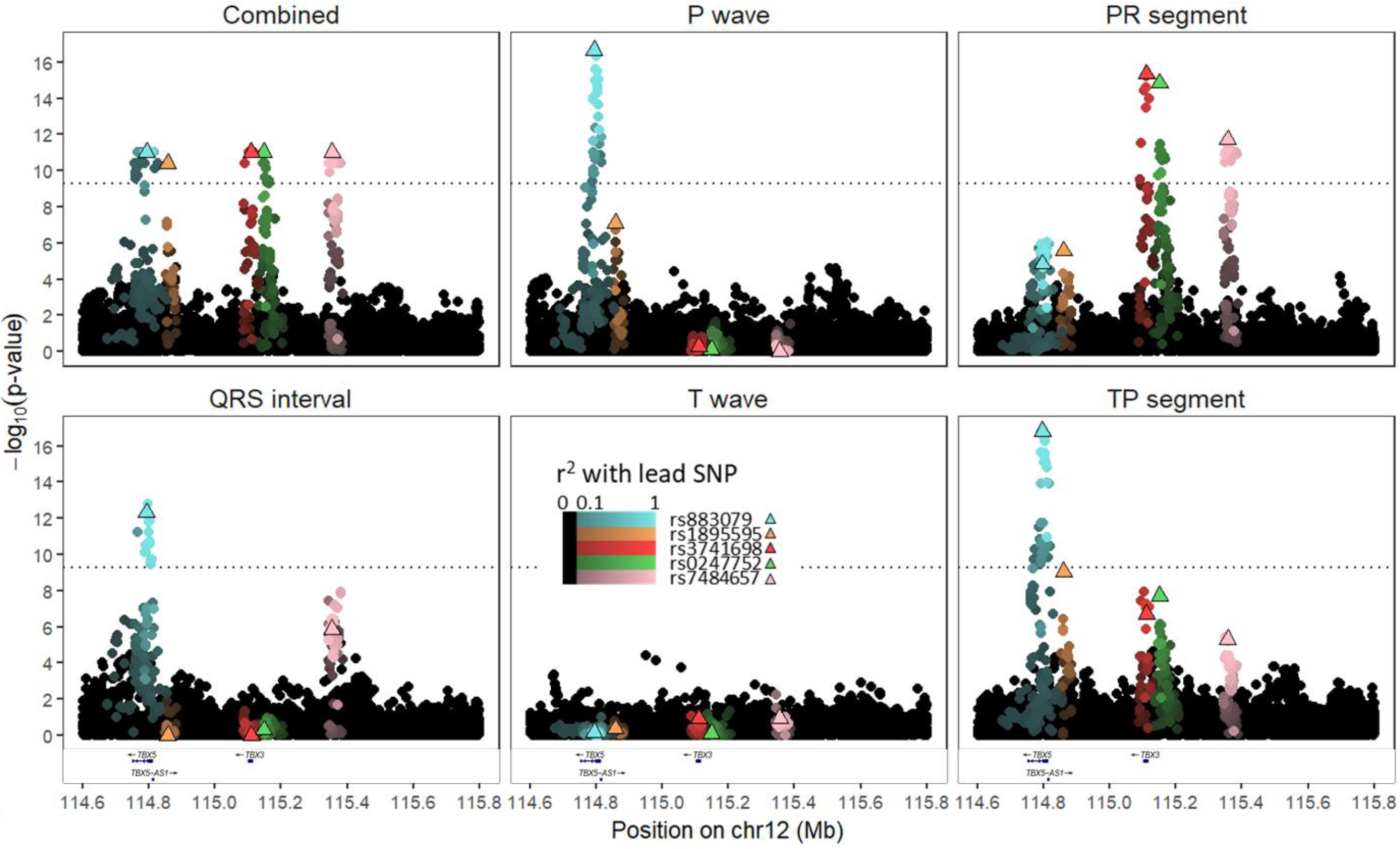
Regional SNP associations and linkage disequilibrium at four independent signals near *TBX5* among 34,668 participants with electrocardiographic data in the Population Architectures using Genomics and Epidemiology (PAGE) study and Multi-Ethnic Study of Atherosclerosis. Lighter colors indicate greater linkage disequilibrium with lead SNPs, and black markers denote SNPs not in LD (r^2^ < 0.1) with any of the four lead SNPs. Combined phenotype p-values are truncated at 1×10^−11^.

## DISCUSSION

Using cardiac conduction and repolarization traits as an example, we examined the extent to which combined multi-ethnic GWAS analyses of carefully selected phenotypes that map to well-defined biology improved detection and characterization of loci. We identified six novel loci, five of which were detected only when examining the more precisely defined phenotypes, and a sixth locus that was specific to African ancestral populations. We also showed how leveraging evidence of a shared genetic architecture aided the characterization of known loci, particularly when loci harbored multiple independent signals that differed by trait. In this mega-GWAS era involving predominantly European ancestral populations, this study, conducted in a population one-third the size of the largest published ECG trait GWAS (13,14), underscores the merits of prioritizing diversity and phenotype measurement.

Of the three GWAS challenges we examined, our deliberate selection of phenotype measures mapping to well-defined biology largely drove locus discovery, challenging current trends in GWAS that emphasize increased sample size. The growing scale of GWAS, which today can surpass one million participants(15), has resulted in the prioritization of commonly available traits (e.g., body mass index) over traits that more precisely capture underlying biology (e.g., direct measures of body fat(16)). In our case, composite ECG traits PR interval and QT interval have been most commonly interrogated by GWAS. However, these traits represent aggregates of physiologically distinct mechanisms, which may obscure loci with effects localized to, or inconsistent across, individual contiguous traits. This phenomenon was illustrated by the *PITX2* locus, a locus associated with atrial fibrillation.(17) Because *PITX2* lead SNP rs13143308 had opposing associations with the contiguous P wave and PR segment, a standard approach using the composite PR interval yielded a near-zero effect, obscuring the potential importance of the locus on atrial function regardless of sample size. These results emphasize the need to balance ongoing investments in large-scale genome measurement with use of precision phenotyping, for instance through efforts like the ongoing Precision Medicine Initiative’s *All of Us* Research Program(18).

The six traits we used in our ECG decomposition were motivated by their relations to physiology, and their coherence as an aggregate electrophysiologic phenotype. While an important complement to traditional, coarser measures like the PR and QT intervals, our phenotype decomposition approach that identified novel loci and improved characterization of known loci captured but a fraction of the full variation in ECG phenotypes; further phenotypic specificity and additional biologic insight may be offered by GWAS of other ECG traits, including measures of waveform amplitudes, angles, or variability. A complementary approach might select traits that are governed by a shared genetic architecture, such as ion channel function or cardiac remodeling when applied to ECG trait GWAS, potentially assisting efforts to map loci to specific biologic pathways. Further extending combined phenotype ECG trait GWAS to include other phenotypes and traits (e.g. cardiometabolic traits or cardiovascular diseases) also is warranted, given evidence that these traits represent interrelated manifestations of common biologic mechanisms (12) as well as success of prior combined phenotype studies to disentangle complex biology(19). Overall, the question of how to select intermediate traits and integrate such traits with other phenotypic data, including clinical and prognostic information, remains an open question, with best practices that likely will vary across complex traits.

Despite mounting interest in combined phenotype statistical approaches, their merits for novel locus discovery and locus characterization remain largely untested. Here, combined phenotype analysis of contiguous ECG traits did not identify novel loci that eluded traditional univariate analyses, despite the purported ability of combined phenotype methods to increase statistical power. However, preliminary evaluation of *TBX5*, a locus harboring multiple independent signals, suggested that combined phenotype approaches may be particularly informative for fine-mapping. Supporting the extension of combined phenotype methods to fine-mapping are methods that have been specifically developed for this challenge(20), including fastPAINTOR. When compared with single trait fine-mapping, fastPAINTOR reduced the number of SNPs required for follow-up in order to capture 90% of the causal variants, from 23 SNPs per locus using a single trait to 12 SNPs when fine-mapping two traits simultaneously. Other potentially fruitful avenues may extend combined-phenotype studies of ECG traits to include cardiovascular disease.

The lack of diversity in GWAS has long been described(21), but the literature remains dominated by studies on European ancestral populations. As a result, genomics research is confined to a narrow sliver of human genetic diversity, even as the US population becomes more diverse(22). Our deliberate selection of an ancestrally diverse population enabled the identification of a novel *CD36* locus, which was common only in populations of African descent. Lead SNP, rs3211938, had a large effect on QT interval, among the largest effects reported to-date,(13) although winner’s curse may be a concern(13). Variant rs3211938, a ClinVar-indexed missense mutation known to cause CD36 deficiency, encodes a scavenger receptor central to formation and cellular uptake of long-chain fatty acids. Although *CD36* and rs3211938 have been associated with a spectrum of cardiometabolic phenotypes,(23–30) the most intriguing finding is the potential linkage with sudden cardiac arrest (SCA), for which QT interval prolongation increases risk(31). SCA accounts for approximately 10-20% of total mortality in industrial countries(32), and several decades of research have suggested a contributory role of impaired fatty-acid uptake in cardiomyocytes. Although genetic studies of *CD36* and SCA were largely null(33,34), the use of predominantly European ancestral populations constrained evaluation of rs3211938, which is near monomorphic in all populations except those of African descent. Overall, these results highlight the potential for racially/ethnically diverse studies to provide novel biological insights, beyond the reach of predominantly European ancestral populations.

Limitations of our study point to several promising directions for future work. First, we lacked a replication cohort, reflecting the rarity of multi-ethnic studies with high-resolution ECGs from which to derive the six contiguous ECG traits. However, this study is the largest multi-ethnic GWAS of ECGs to-date, with excellent statistical power, and we identified loci that are biologically plausible. Second, we limited our evaluation to common variants, although previous studies have demonstrated the utility of interrogating rare variants, particularly in the context of multi-ethnic studies(35,36). Our focus on common variants reflects the current limitations of combined phenotype methods for interrogating rare variants in complex samples or with summary data. Widespread interest in this approach suggests that this gap may be closed soon. Further, while this study helps address the lack of diversity in ECG trait GWAS, the small number of Chinese American participants limited our ability identify loci that were common only in populations of East Asian descent. Future efforts that further expand population racial/ethnic diversity represents an important next both for cardiac conduction studies and GWAS more broadly. Finally, we did not perform in-depth fine-mapping, although approaches that leverage multiple phenotypes have been described(20). Identification of allelic heterogeneity provides a clear impetus for future studies that leverage evidence of a shared genetic effects to disentangle the genetic architecture underlying ECG traits and, more broadly, complex traits.

This study illustrates three strategies to improve the efficiency of locus discovery. Of these, our findings emphasize the importance of carefully constructed phenotypes and of ancestral diversity for novel locus identification. In contrast, combined phenotype methods did not enable identification of novel loci unapparent using traditional approaches, although combined phenotype studies did inform characterization of known loci. As researchers contemplate the next generation of genomics studies, increased phenotype resolution and ancestral diversity will be crucial to understanding the ever-expanding “phenome,” while ensuring equitable access to precision medicine.

## MATERIAL AND METHODS

### Study population

#### Data sources

The multi-ethnic PAGE study(6) is a consortium funded by the National Human Genome Research Institute (NHGRI) to examine the genetic underpinnings of common complex diseases and traits in ancestrally-diverse populations (online supplement). PAGE data used in this study included African American, European American, and Hispanic/Latino participants enrolled in the Women’s Health Initiative (WHI)(37) Clinical Trial, and in the population-based Atherosclerosis Risk in Communities Study (ARIC)(38), and Hispanic Community Health Study/Study of Latinos (HCHS/SOL)(39). Data for African American, Chinese American, European American, and Hispanic/Latino populations also were contributed by the population-based Multi-Ethnic Study of Atherosclerosis (MESA)(40) (online supplement).

### Electrocardiography

Ten-second, resting, standard 12-lead ECGs were collected following standard protocols (online supplement). To enable direct comparison with the published literature, we used exclusion criteria from published GWAS of QT interval, QRS duration, PR interval, and P wave. Poor-quality ECGs and ECGs exhibiting severe abnormalities were excluded (Table S1, S2). Also excluded were participants with prevalent heart failure or coronary heart disease, and participants taking any class I or III anti-arrhythmic medication.

We temporally decomposed the ECG waveform into six contiguous ECG traits, each reflecting distinct physiological processes: P wave, PR segment, QRS interval, ST segment, T wave, and TP segment (Figure 4). Three of the six contiguous traits were measured directly from maximum readings across all 12 ECG leads (P wave, QRS interval, T wave), and were used to construct the remaining three traits, as detailed in supplementary material. We also evaluated two composite ECG traits that are widely available and have been examined in numerous GWAS analyses: the QT interval(13) and the PR interval(14). The PR interval combines the contiguous P wave and PR segment, reflecting conduction through the atria and the AV-node. The QT interval spans the contiguous QRS interval, ST segment, and the T wave, representing ventricular activity.

**Figure 4.**
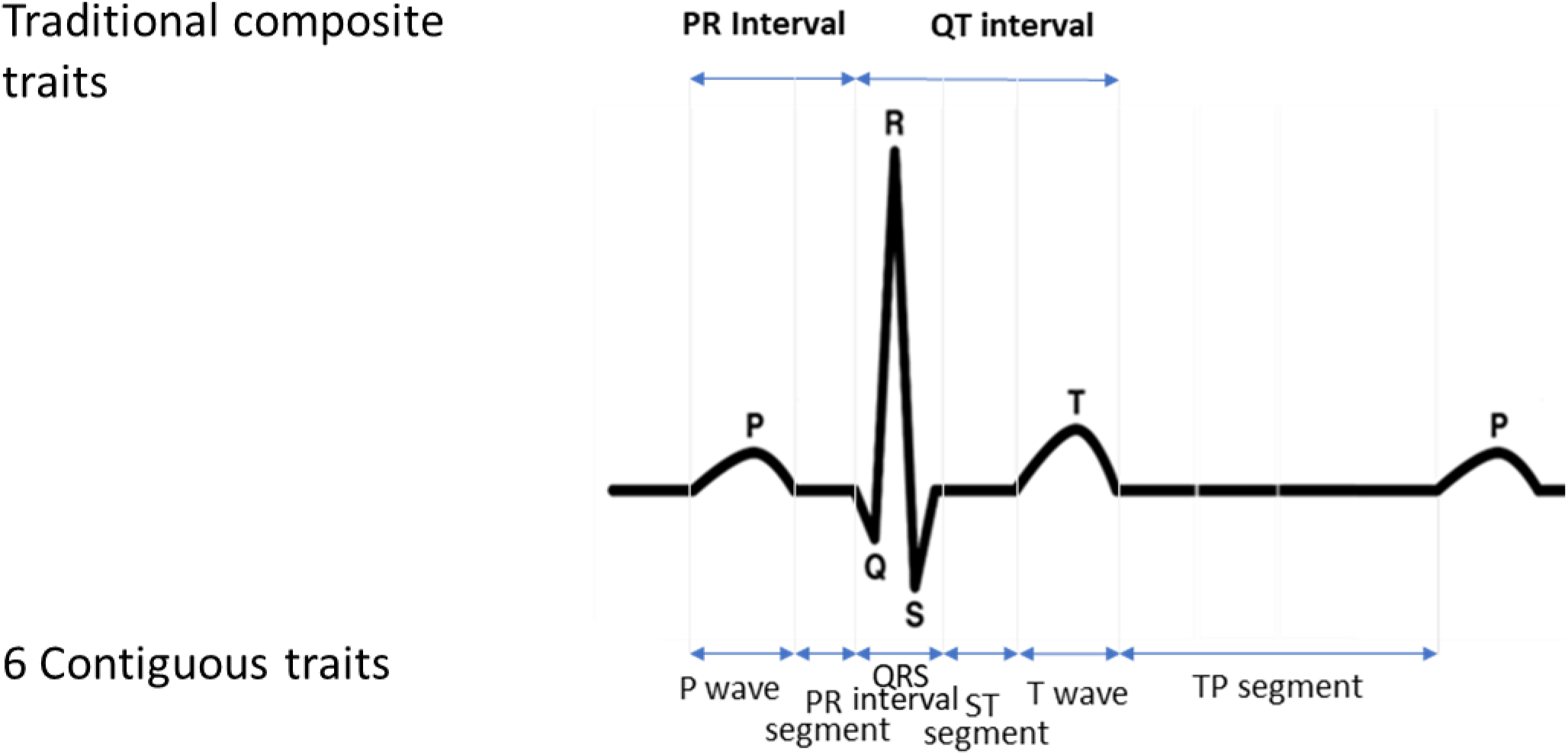
Illustration of the six contiguous (P wave, PR segment, QRS interval, ST segment, and TP segment) and two composite (QT interval and PR interval) ECG traits.

### Genotyping and Imputation

Imputation was performed across studies, using the 1000 Genomes Project phase 3 reference panel (Table S3). In addition to study-specific quality control protocols, we excluded SNPs meeting any of the following criteria: minor allele frequency (MAF) <1%; imputation quality score < 0.3; or effective sample size < 30, calculated as 2 × *MAF*(1 − *MAF*) × *N* × *Q*, where *Q* is imputation quality and *N* is study size.

### Statistical Methods

#### Overview

Statistical analyses were carried out in two stages. First, genome-wide univariate associations for each of six contiguous ECG traits were calculated in twelve substudies defined by genotyping platform, race/ethnicity, and study (Table S3). The pooled PAGE sample of Hispanic/Latino and African American participants from the HCHS/SOL and WHI genotyped together(41) on the Multi-Ethnic Genotyping Array (MEGA) array was retained as a single substudy. The remaining 11 substudies were defined by ancestry (African American, Chinese American, and European American) and study (ARIC, MESA, and WHI). Substudy-specific univariate results for the six contiguous ECG traits were then combined via trait-specific inverse-variance meta-analysis(42). Next, meta-analyzed contiguous ECG trait-specific univariate associations were evaluated in aggregate using the adaptive sum of powered scores method(43) (aSPU, described below), yielding a combined-trait p-value for each SNP. All univariate and combined-phenotype analyses were performed in the multi-ethnic population as demonstrated previously in the PAGE study(44), and, in sensitivity analyses, by self-identified race/ethnicity.

#### Univariate associations

Univariate associations for the six contiguous ECG traits and the two composite ECG traits were estimated assuming an additive genetic model of inheritance and adjusting for linear effects of age at study exam, sex, study site or region, inverse heart rate, and ancestral principal components(45). The pooled PAGE sample was analyzed using generalized estimating equations allowing correlated errors among households sharing first or second-degree relatives, and independent error distributions by self-reported ancestry group(46). Linear regression was used for the other 11 analytic groups. Models were implemented using SUGEN (46) (ARIC, pooled PAGE sample, WHI) and SNPTest (MESA). For each continuous and composite ECG trait, results were combined across substudies using inverse-variance weighted, genomic inflation-corrected meta-analysis(42). SNP effect heterogeneity across substudies and, in sensitivity analyses, among self-reported race/ethnicity was measured with the Cochran’s Q test. For each ECG trait, SNP meta-analysis p-values were assessed by calculating genomic inflation factors (λ) and plotting the expected distribution against observed results.

#### Combined phenotype analyses

To evaluate evidence of shared genetic effects across all six contiguous ECG traits, meta-analyzed univariate results were combined with aSPU(43). The aSPU procedure was selected based on its accommodation of effect direction across traits, low type I error rate in simulation studies when compared with other available procedures (data not shown), and its scalability to 1000 genomes imputed data. We implemented aSPU using Julia 1.0 to optimize efficiency, and made the code freely available as a Julia package (https://github.com/kaskarn/JaSPU).

aSPU uses meta-analyzed univariate summary z-scores, calculated for each SNP across all *k* traits. Briefly, the procedure estimates Σ, the *k* × *k* correlation of null z-scores across univariate results and draws 10^11^ Monte-Carlo samples from the multivariate *N*_*k*_ (0, Σ) distribution. For each SNP *j*, the results for all *k* traits *Z*_*j*1_, *Z*_*j*2_, …, *Z*_*jk*_ are used to form the sequence of powered scores: 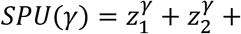 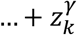, where γ = 0, 1, …, 8, ∞. This sequence is compared to values produced by all 10^11^ draws from the simulated null distribution, and a Monte-Carlo p-value is derived for the best-powered SPU(γ) to calculate an overall SNP-p-value (*p*_*aSPU*_), ranging between 1/(1+10^11^) and 1. The test is adaptive in the sense that the γ yielding the maximal SPU(γ) can vary across SNPs, so that the test maintains power against a number of different alternative hypotheses.

### Reporting

Multi-ethnic combined phenotype results were presented as the primary findings, employing Bonferroni correction assuming 10M SNP tests (i.e. p_aSPU_<5×10^−9^). Ancestry-specific and ECG-trait specific analyses were performed to aid interpretation of results and are considered sensitivity analyses.

Previously-reported SNPs were identified through review of the NHGRI-EBI GWAS Catalog(12) as of January 22, 2019 (Table S6), for the following interval scale ECG phenotypes: PR interval, PR segment, QRS interval, P wave, QT interval, and T wave, supplemented by a literature review to identify publications not indexed by the NHGRI-EBI GWAS Catalog(14,44,47). There were no published GWAS analyses of ST segment or TP segment durations.

We defined a locus to encompass all SNPs within 500 kb of, and in linkage-disequilibrium (LD) with, the lead SNP (multi-ethnic r^2^ > 0.2 in the unrelated PAGE study participants). SNPs > 500 kb away from all previously reported ECG trait loci were considered novel. Locus-specific lead SNPs were identified as the SNP with the lowest paSPU. Tied lowest paSPU were resolved using the lowest locus-specific univariate p-value across contiguous ECG traits. Complete summary level results from all analyses are made available through dbGaP (phs000356).

### Ethics Statement

All data were analyzed anonymously, and this study was exempted from review under 45 CFR 46.101(b) by the University of North Carolina institutional review board (IRB #18-1764).

## SUPPORTING INFORMATION

Supporting material and methods, tables (S1-9), and figures (S1-2) are included in two additional files

**Figure S1. Quantile-Quantile plots for trans-ancestry meta-analyzed associations of SNPs with each of six ECG traits (decomposed ECG phenotype)** in n=34,668 participants from the Population Architectures using Genomics and Epidemiology (PAGE) study and the Multi-Ethnic Study of Atherosclerosis (MESA). Black markers and lambda values represent to p-values for all SNPs that passed quality control. Blue markers represent the subset of SNPs >500kb from any previously-reported ECG lead SNP.

**Figure S2 Manhattan plots for univariate, trans-ethnic ECG trait GWAS** in 34,668 participants from the Population Architectures using Genomics and the and Multi-Ethnic Study of Atherosclerosis (MESA)‥ Genome-wide (p_univariate_ < 5×10^−9^) significant loci within 500kb of previously-reported ECG GWAS results are shown in yellow; genome-wide (p_univariate_ < 5×10^−9^) significant loci >500 kb from previously reported ECG loci are shown in blue.

**Table S1.** Exclusion criteria for genome-wide study of all evaluated electrocardiographic traits.

**Table S2**. Participant exclusions (cumulative), by study and ancestry group

**Table S3**. Genotyping, imputation, and quality control by participating study

**Table S4. Participant characteristics by study of origin, and ancestry group**, among N=34,668 participants from the Population Architecture using Genomics and Epidemiology (PAGE) study and the Multiethnic Study of Atherosclerosis (MESA).

**Table S5. Partial correlations between six contiguous traits decomposing the ECG**, in n=10,618 eligible Hispanic Community Health Study/Study of Latinos participants. Partial correlations were adjusted for RR interval, gender, study site, and ancestry principal components.

**Table S6. Published genome-wide association studies of electrocardiographic traits**, indexed on the NHGRI GWAS catalog (Sept. 30, 2018).

**Table S7. Previously-reported associations of SNPs with temporal electrocardiographic traits**, and corresponding results in N=34,668 participants in the Population Architecture using Genomics and Epidemiology (PAGE) study and the Multiethnic Study of Atherosclerosis (MESA).

**Table S8 Trait-specific direction of effects, and meta-analysis heterogeneity p-values for univariate associations (punivariate < 5×10-9) of loci discovered in combined-phenotype analyses of six electrocardiographic traits (paSPU < 5×10-9),** in 34,668 participations from the Population Architecture usingGenomics and Epidemiology study (PAGE) and Multi-Ethnic Study of Atherosclerosis (MESA).

**Table S9 Trait-specific trans-ethnic meta-analyzed effect estimates and standard errors for univariate associations (punivariate < 5×10-9) of loci discovered in combined-phenotype analyses of six electrocardiographic traits (paSPU < 5×10-9)**, in 34,668 participations from the Population Architecture using Genomics and Epidemiology study (PAGE) and the Multi-Ethnic Study of Atherosclerosis (MESA).

**Table S10** Imputation quality scores of all novel variants reported in this study.

## ACKNOWLEDGMENTS

The contents of this paper are solely the responsibility of the authors and do not necessarily represent the official views of the National Institutes of Health.

The PAGE consortium thanks the staff and participants of all PAGE studies for their important contributions. The complete list of PAGE members can be found at http://www.pagestudy.org.

The Stanford Global Reference Panel was created by Stanford - contributed samples and comprises multiple datasets from multiple researchers across the world designed to provide a resource for any researchers interested in diverse population data on the Multi-Ethnic Global Array (MEGA). The authors thank the researchers and research participants who made this dataset available to the community.

The authors thank the WHI investigators and staff for their dedication, and the study participants for making the program possible. A full listing of WHI investigators can be found at: http://www.whiscience.org/publications/WHI_investigators_shortlist.pdf.

The authors thank the staff and participants of the ARIC study for their important contributions. More detail about the ARIC study may be found at: https://sites.cscc.unc.edu/aric/.

Genotyping for MESA was performed at Affymetrix (Santa Clara, California, USA) and the Broad Institute of Harvard and MIT (Boston, Massachusetts, USA) using the Affymetrix Genome-Wide Human SNP Array 6.0. We also thank the other investigators, the staff and the participants of MESA for their valuable contributions. A full list of participating MESA investigators and institutions can be found online (http://www.mesa-nhlbi.org).

