## Supplemental materials for "Multi-ethnic genome-wide association study of decomposed cardioelectric phenotypes illustrates strategies to identify and characterize evidence of shared genetic effects for complex traits"

### SUPPLEMENTAL MATERIAL AND METHODS

### Participating studies

#### The Atherosclerosis Risk in Communities Study (ARIC)

The Atherosclerosis Risk in Communities Study (ARIC) is a longstanding study sponsored by the National Heart, Lung and Blood Institute to investigate cardiovascular disease causes, and clinical outcomes. Recruitment of predominantly Caucasian and African-American descent participants occurred between 1987-1989 from four communities in the United States (Forsyth County NC, Jackson MS, suburban Minneapolis MN, and Washington County MD). In total, 15,792 participants were enrolled into the cohort, and received standardized physical examinations and interviewer-administered questionnaires at study baseline (1987-1989), from which the electrocardiographic data used in this study were obtained. Participants were genotyped on the Affymetric GeneChip SNP 6.0 Array.

#### Hispanic Community Health Study / Study of Latinos (HCHS/SOL)

The HCHS/SOL is a multicenter, community-based cohort study of U.S. Hispanic/Latinos. Goals of the study are to examine the prevalence of and risk factors for several disorders including heart, lung, blood, and kidney phenotypes^1^. HCHS/SOL investigators sampled 16,415 males and females aged 18-74 years at baseline from four study communities: the Bronx, NY, Chicago, IL, Miami, FL, and San Diego, CA. HCHS/SOL recruitment centers were selected so that the study would include at least 2,000 participants in each of the following designations: Mexican, Puerto Rican, Dominican, Cuban, and Central and South American. The two-stage sampling design selected households within census block groups. Households with Hispanic/Latino surnames and individuals over 45 years of age were oversampled to achieve increased representation of Hispanic/Latino individuals with a uniform age distribution. For this study, we used ECG data collected at the baseline exam (2008-11) in n=12,803 participants genotyped on the multi-ethnic genotyping array (MEGA).

#### The Multi-Ethnic Study of Atherosclerosis (MESA)

MESA was initiated in 2000 to investigate subclinical cardiovascular disease and the risk factors that predict progression to clinically overt cardiovascular disease^2^ in a multi-ethnic U.S. population free of clinically-recognized cardiovascular disease at study baseline. The population-based cohort included 6,814 men and women) of European American (38%), African American (28%), Hispanic (22%) and Chinese American (12%) descent, 45–84 years of age from six field centers (Winston-Salem, NC; St. Paul, MN; Chicago, IL; Los Angeles, CA; New York, NY; Baltimore, MD. For this study, ECG measures were collected during the baseline visit, with the exception of T wave, which was collected at visit 5. Sensitivity analyses verified that the inclusion of visit 5 data did not meaningfully influence results of combined-phenotype analyses using the associated sum of powered scores test (aSPU).

#### The Women’s Health Initiative (WHI)

The Women’s Health Initiative was initiated by the National Institutes of Health in 1991. It consists of three clinical trials (CT), and an observational study (OS), totaling 168,132 women distributed across the United States. The present analyses included European American participants from the observational study, and controls from the clinical trials genotyped on the Illumina Omni-Quad Array, Affymetrix Gene Titan, and the Axiom Genome-Wide Human CEU I Array: GECCO (N=930 WHI-CT controls; N=3720 WHI-OS); the Hip Fracture Study (HipFX), the Modification of PM-Mediated Arrhythmogenesis in Populations study (MOPMAP, N=9,014 WHI-CT controls); the Women's Health Initiative Memory Study (WHIMS, N=5,740 WHI-CT cases and controls); and the Genome-wide Association Research Network into Effects of Treatment (GARNET, N=4,883, WHI-CT controls). African American and Hispanic/Latino women genotyped using the MEGA array by the PAGE study were also included, and were analyzed jointly with MEGA array data from the HCHS/SOL study (above). Sensitivity analyses demonstrated that adjustment for clinical trial participation did not alter study results.

### Electrocardiographic (ECG) traits

In all four studies, standard resting, supine or semi-recumbent 12-lead ECGs were administered during a baseline visit, and transmitted to the Wake Forest School of Medicine Epidemiology Cardiology Research Center (EPICARE) for automatic processing using contemporary versions of General Electric’s 12SL Marquette computerized analysis program (GE, Milwaukee, WI).

We used maximum measurements across all twelve leads for P-wave, QRS interval and T wave durations, and derived the three isoelectric segments from additional data on QT interval, PR interval, and RR duration calculated from inverse-heartrate. Respectively, PR, ST and TP segment durations were obtained by subtracting the P-wave from the PR interval, the QRS interval and T-wave durations from the QT interval, and the QT and PR-intervals from the RR-interval. Since temporal measurements corresponded to maximum values across leads, negative PR-segment or ST-segment values were calculated in a small number of participants.

This decomposition into six contiguous traits was used to map ECG traits to distinct phases of the average heartbeat, and is summarized below.

| **Contiguous ECG trait** | **Coverage** | **Meaning** | **Measured or calculated** |
| --- | --- | --- | --- |
| P wave | Start of P wave to end of P wave | Atrial depolarization | Measured on tracings |
| PR segment | End of P wave to start of Q wave | Conduction through the atrioventricular node and bundle of His | Calculated as PR interval–P wave |
| QRS interval | Start of Q wave to end of S wave | Ventricular depolarization | Measured on tracings |
| ST segment | End of S wave to start of T wave | Electrically silent; ventricles stay contracted | Calculated as QT interval – QRS – T wave |
| T wave | Start of T wave to end of T wave | Ventricular repolarization | Measured on tracings |
| TP segment | End of T wave to start of P wave | Electrically silent; refractory stage preceding the next heartbeat | Calculated as RR – QT – PR interval |

**SUPPLEMENTAL FIGURES**

**Figure S1** **Quantile-Quantile plots for trans-ancestry meta-analyzed associations of SNPs with each of six ECG traits (decomposed ECG phenotype)** in n=34,668 participants from the Population Architectures using Genomics and Epidemiology (PAGE) study and the Multi-Ethnic Study of Atherosclerosis (MESA). Black markers and lambda values represent to p-values for all SNPs that passed quality control. Blue markers represent the subset of SNPs >500kb from any previously-reported ECG lead SNP.

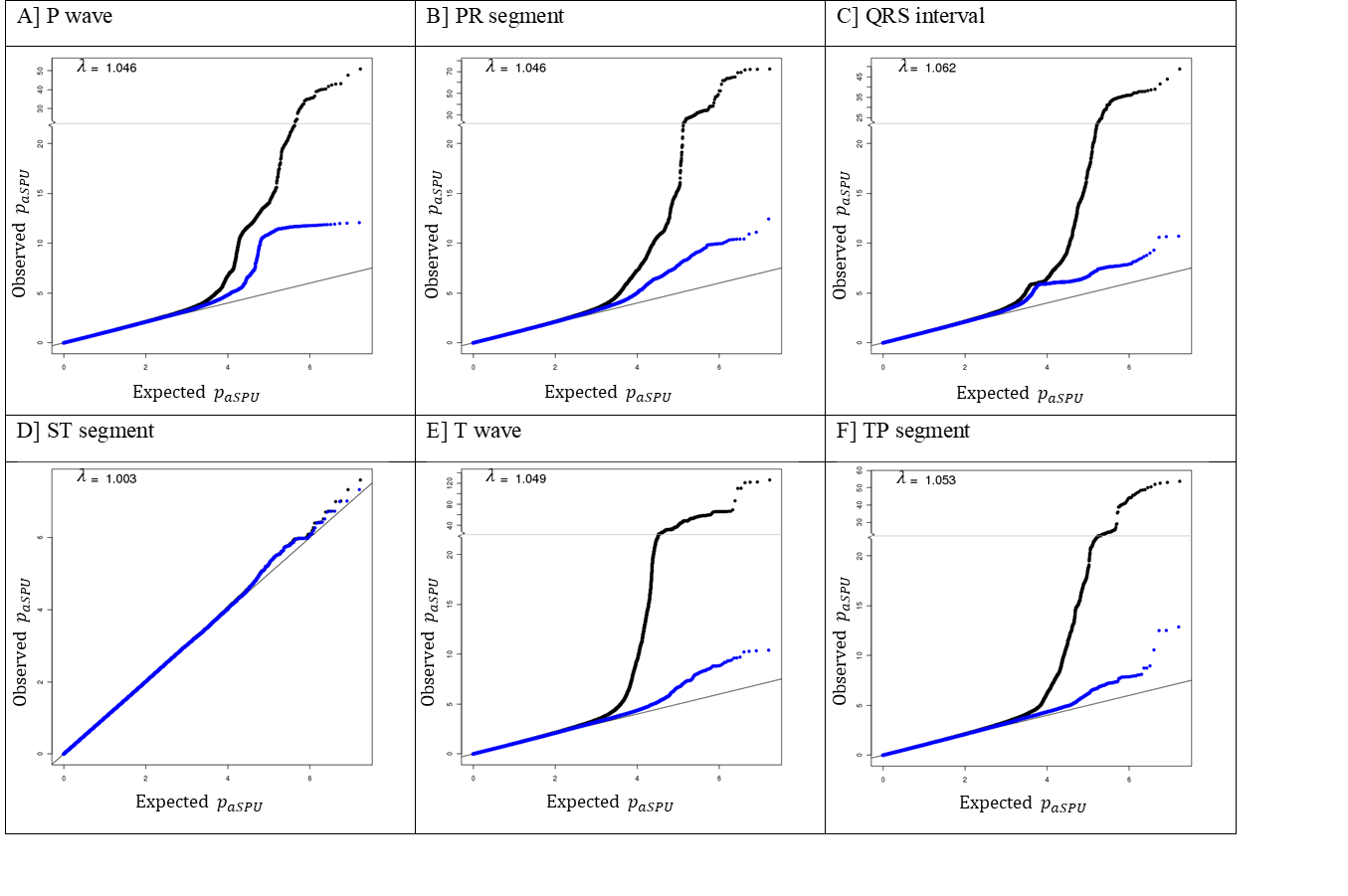

**Figure S2 Manhattan plots for univariate, trans-ethnic ECG trait GWAS** in 34,668 participants from the Population Architectures using Genomics and the and Multi-Ethnic Study of Atherosclerosis (MESA).. Genome-wide (p_univariate_ < 5×10^-9^) significant loci within 500kb of previously-reported ECG GWAS results are shown in yellow; genome-wide (p_univariate_ < 5×10^-9^) significant loci >500 kb from previously reported ECG loci are shown in blue.

A] P wave

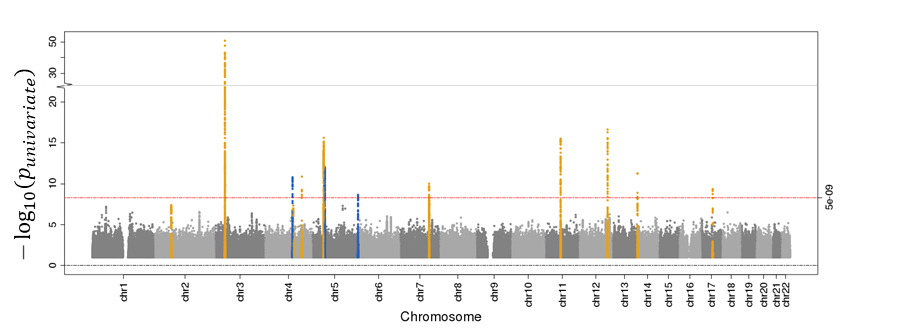

B] PR segment

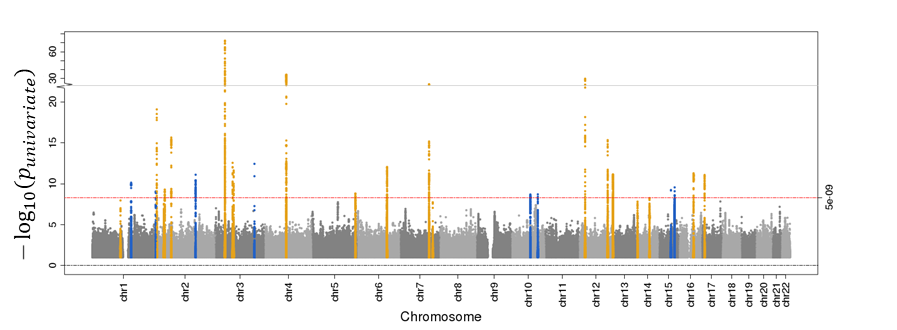

C] QRS interval

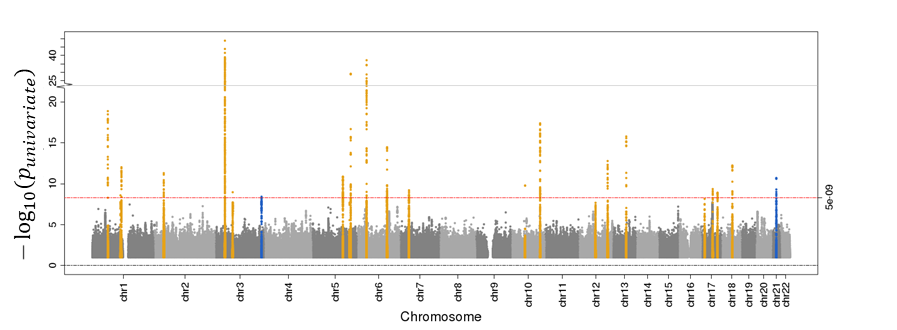

D] ST segment

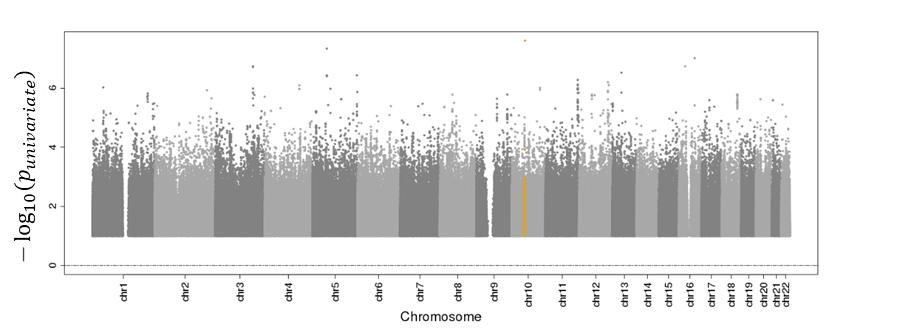

E] T wave

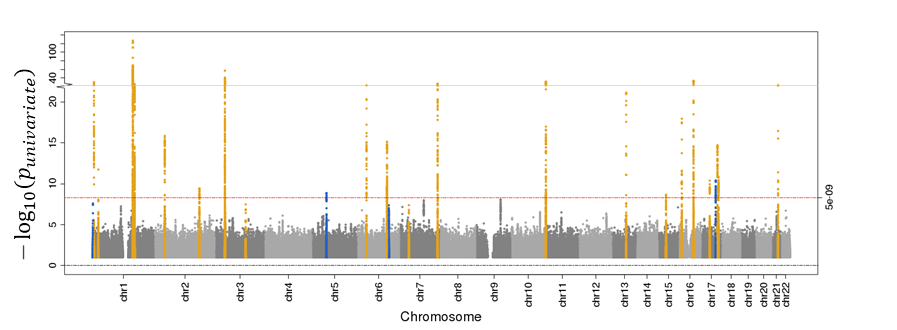

F] TP segment

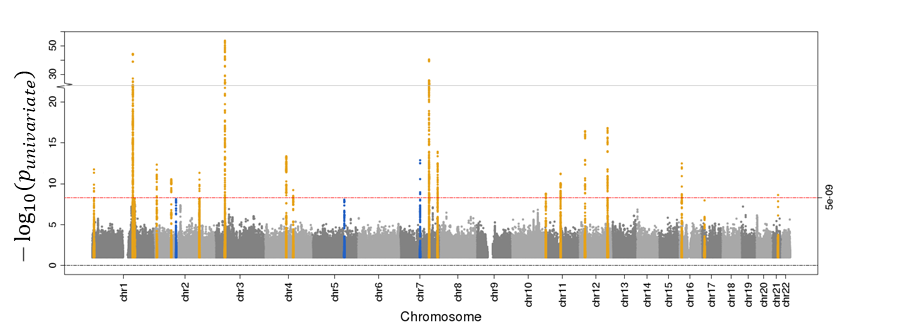

**SUPPLEMENTAL TABLES**

| **Table S1.** **Exclusion criteria for genome-wide study of all evaluated electrocardiographic traits.** | |
| --- | --- |
| Exclusion criterion | Description |
| Low ECG quality | ECG quality rated as 5 |
| Pacemaker | Any artificial pacemaker |
| Prevalent CHD | Prevalent MI, or non-angina CHD |
| Bundle-branch block | Major ventricular conduction delay: MCs 7.1.1, 7.2.1, 7.4, or 7.8 |
| Wolf-Parkinson-White (WPW) pattern | Definite evidence of WPW pattern: MC 7.4.1 |
| Prevalent atrial fibrillation or flutter | Persistent atrial fibrillation: MCs 8.3.1, 7.3.2 |
| Atrioventricular block | Second or third degree atrioventricular (AV) block, or Mobitz II pattern. MCs 6.1, 6.2.1, 6.2.2 |
| Ectopic beat | Study-specific thresholds |
| Antiarrhythmics use | Class I and III antiarrhythmic use, assessed at baseline drug inventories |
| Prevalent heart failure | HCHS/SOL and WHI: self-reported doctor-diagnosed heart failure ARIC: prevalent CHD, or edema, paroxysmal nocturnal dyspnea, angina, rales, or atrial fibrillation (Gothenburg criterion)  MESA: self-reported doctor-diagnosed heart failure |
| Outlier ECG values | > 4 SD from ancestry and study average, for any of: P wave, PR segment, QRS interval, ST segment, T wave, or TP segment duration |

ARIC: Atherosclerosis Risk in Communities Study, CHD: Coronary Heart Disease, ECG: Electrocardiogram, HCHS/SOL: Hispanic Community Health Study/Study of Latinos, MC: Minnesota Code(s), MESA: Multi-Ethnic Study of Atherosclerosis, MI: Myocardial Infarction, SD: Standard Deviation, WHI: Women’s Health Initiative

| **Table S2**. **Participant exclusions (cumulative), by study and ancestry group** | | | | | | | | | | |
| --- | --- | --- | --- | --- | --- | --- | --- | --- | --- | --- |
|  | **ARIC** | | **MESA** |  |  |  | **HCHS/SOL*** | **WHI** |  |  |
| Exclusion criterion | AA | EA | EA | AA | HIS | CHN | HIS | AA* | EA | HIS |
| Low ECG quality | 12 (12) | 56 (56) | 15(15) | 14(14) | 11(11) | 13(13) | 102 (102) | 53 (53) | 87 (87) | 17 (17) |
| Pacemaker | 3 (15) | 8 (64) | 0(15) | 0(14) | 0(11) | 0(13) | 13 (115) | 7 (53) | 9 (87) | 2 (18) |
| Prevalent CHD | 115 (128) | 452 (508) | 0(15) | 0(14) | 0(11) | 0(13) | 630 (727) | 229 (281) | 473 (558) | 50 (68) |
| Bundle-branch block | 72 (194) | 242 (706) | 13(28) | 5(19) | 4(15) | 0(13) | 265 (947) | 111 (387) | 322 (849) | 30 (98) |
| Wolf-Parkinson-White pattern | 2 (195) | 3 (709) | 0(28) | 0(19) | 1(16) | 0(13) | 21 (953) | 0 (387) | 0 (849) | 0 (98) |
| Prevalent atrial fibrillation | 0 (195) | 0 (709) | 0(28) | 0(19) | 0(16) | 1(14) | 0 (953) | 7 (393) | 21 (865) | 1 (98) |
| Atrioventricular block | 3 (195) | 8 (709) | 0(28) | 0(19) | 0(16) | 0(14) | 16 (954) | 1 (393) | 2 (867) | 0 (98) |
| Ventricular ectopy | 1 (195) | 18 (727) | 49(77) | 31(50) | 35(51) | 5(19) | 35 (987) | 28 (414) | 115 (970) | 9 (106) |
| Antiarrhythmics use | 16 (202) | 67 (758) | 382(459) | 454(504) | 246(297) | 134(153) | 12 (996) | 14 (423) | 32 (994) | 0 (106) |
| Prevalent heart failure | 186 (347) | 334 (1008) | 0(459) | 0(504) | 0(297) | 0(153) | 219 (1103) | 59 (453) | 59 (1021) | 12 (117) |
| Outlier ECG values | 29 (376) | 61 (1069) | 3(462) | 4(508) | 2(299) | 0(153) | 52 (1155) | 26 (479) | 95 (1116) | 9 (126) |
| Remaining (%): | 2,387 (86%) | 8,046 (88%) | 2065 (82%) | 1168(70%) | 1172(81%) | 622 (80%) | 10,618 (90%) | 3,759 (89%) | 8,834 (89%) | 1,573 (93%) |

AA: African American, CHN: Chinese-American, EA: European-American, HIS: Hispanic/Latinos

ARIC: Atherosclerosis Risk in Communities Study, CHD: Coronary Heart Disease, ECG: Electrocardiogram, HCHS/SOL: Hispanic Community Health Study/Study of Latinos, MESA: Multi-Ethnic Study of Atherosclerosis, SD: Standard Deviation, WHI: Women’s Health Initiative

*HCHS/SOL, WHI AA and WHI HIS were combined in analyses

| **Table S3**. **Genotyping, imputation, and quality control by participating study** | | | | | | | | |
| --- | --- | --- | --- | --- | --- | --- | --- | --- |
|  | ARIC | GARNET | GECCO | HIPFX | MOPMAP | WHIMS | PAGE* | MESA |
| **Genotyping Platform** | Affymetrix GeneChip SNP Array 6.0 | Illumina Human Omni1-Quad v1-0 B | Illumina 610 and Cytochip 370K | Illumina 50K and 610K | Affymetrix Gene Titan, Axiom Genome-Wide, Human CEU I Array Plate | Human OmniExpress Exome-8v1_B Genome-Wide Human | Infinium Expanded Multi-Ethnic Genotyping Array | Affymetrix  GeneChip  SNP Array  6.0 |
| **QC Filters** |  |  |  |  |  |  |  |  |
| Sample call rate | 90% | 98% | 98% | 98% | 90% | 98% | 98% | 90% |
| HWE threshold | p>10^-6^ | | | | | | | |
| **Imputation Software** | MaCH v1.0.16 | BEAGLE | BEAGLE | BEAGLE | BEAGLE | BEAGLE | IMPUTE version 2.3.2 | IMPUTE  version 2.3.2 |
| **Reference Panel** | 1000 Genome phase 3 v 5 | | | | | | |  |
| **Build** | Genome Reference Consortium Human Build 37 | | | | | | | |
| **dbSNP version** | 150 | | | | | | | |

ARIC: Atherosclerosis Risk in Communities Study, MESA: Multi-Ethnic Study of Atherosclerosis, PAGE: Population Architecture using Genetic Epidemiology study, WHI: Women’s Health Initiative

* Includes African-American and Hispanic participants from the Women’s Health Initiative, and all participants genotyped in the Hispanic Community Study/Study of Latinos.

| **Table S4. Participant characteristics by study of origin, and ancestry group**, among N=34,668 participants from the Population  Architecture using Genomics and Epidemiology (PAGE) study and the Multiethnic Study of Atherosclerosis (MESA). | | | | | | | | | | |
| --- | --- | --- | --- | --- | --- | --- | --- | --- | --- | --- |
|  | **ARIC** | | **MESA** | | | | **WHI** | | | **HCHS/SOL** |
|  | AA | EUR | EA | AA | HIS | CHN | AA | EUR | HIS | HIS |
| N (% Female) | 2,387 (63%) | 8,046 (55%) | 1085 (53%) | 621 (53%) | 591 (50%) | 315 (51%) | 2,400 (100%) | 8,742 (100%) | 1,540 (100%) | 10,618 (60%) |
| Age, years | 53.2 (5.7) | 54.0 (5.6) | 61.5 (10.1) | 60.9 (10.3) | 60.3 (10.3) | 61.1 (10.3) | 59.1 (6.2) | 66.1 (6.6) | 59.8 (6.4) | 45.3 (13.7) |
| P wave, ms | 111.7 (11.6) | 106.4 (11.8) | 104.4 (13.2) | 107.8(13.3) | 105.1 (12.1) | 101.5 (11.8) | 110.6 (11.3) | 107.7 (11.9) | 106.4 (11.5) | 107.0 (11.6) |
| PR segment, ms | 59.3 (23.4) | 53.1 (19.7) | 58.3 (23.7) | 61.4 (27.7) | 56.1 (21.4) | 62.3 (21.2) | 56.1 (21.6) | 53.0 (21.3) | 49.4 (19.0) | 49.3 (19.7) |
| QRS complex, ms | 90.0 (9.8) | 91.0 (9.5) | 92.6 (9.4) | 90.4 (10.0) | 90.7 (9.8) | 89.3 (9.4) | 85.1 (8.9) | 86.0 (8.4) | 85.4 (8.4) | 90.8 (9.4) |
| ST segment, ms | 116.3 (18.3) | 117.6 (15.8) | 106.5 (26.3) | 110.0 (28.5) | 106.2 (25.6) | 103.5 (25.0) | 106.6 (22.3) | 107.5 (19.9) | 107.7 (20.2) | 118.4 (17.9) |
| T wave, ms | 201.6 (19.8) | 199.0 (18.2) | 211.7 (22.2) | 209.0 (24.9) | 210.8 (24.4) | 216.2 (24.3) | 208.3 (19.7) | 207.5 (18.1) | 208.4 (18.8) | 206.2 (23.1) |
| TP segment, ms | 345.2 (127.6) | 355.8 (115.6) | 400 (123.9) | 399.7 (130.2) | 397.4 (119.6) | 392.2 (110.8) | 352.3 (120.1) | 365.6 (111.1) | 371.7 (108.1) | 300.3 (126.0) |
| QT interval, ms | 407.9 (29.0) | 407.6 (25.9) | 410.2 (28.5) | 408.8 (31.0) | 407.1 (28.9) | 409.0 (27.4) | 400.1 (31.9) | 401.0 (28.9) | 401.5 (28.9) | 415.4 (28.1) |
| PR interval, ms | 171.0 (26.0) | 159.5 (22.5) | 162.6 (24.7) | 169.2 (28.0) | 161.2 (21.4) | 163.9 (21.5) | 166.7 (24.1) | 160.7 (23.4) | 155.8 (21.0) | 156.3 (20.1) |
| Heartrate, /min | 66.9 (10.7) | 66.6 (9.6) | 63.3 (9.3) | 63.2 (10.1) | 63.7 (9.3) | 63.5 (8.4) | 67.2 (10.7) | 66.4 (9.8) | 66.2 (9.5) | 62.8 (9.4) |
| BMI, kg/m^2^ | 29.5 (5.9) | 26.8 (4.7) | 27.4 (5.0) | 30.0 (5.9) | 29.2 (5.0) | 23.8 (3.3) | 31.5 (6.6) | 28.5 (6.4) | 29.4 (5.8) | 29.6 (5.9) |
| LDL, mg/dL | 138.1 (43.1) | 136.9 (37.6) | 118.6 (30.3) | 118.2 (33.3) | 121.0 (33) | 115.7 (29) | 145.7 (40.5) | 151.4 (35.7) | 143.4 (36.9) | 123.7 (36.6) |
| HDL, mg/dL | 55.6 (17.4) | 51.3 (16.6) | 53.0 (16.0) | 52.5 (15.4) | 47.5 (13.0) | 49.6 (12.6) | 56.4 (13.9) | 54.3 (12.8) | 51.9 (12.6) | 49.3 (13.1) |
| Triglycerides, mg/dL | 112.6 (83.1) | 133.9 (89.9) | 130.6 (92.4) | 103.5 (62.9) | 157.5 (105.0) | 141.3 (84.8) | 112.8 (60.9) | 142.2 (75.6) | 158.5 (82.4) | 137.1 (109.6) |
| Total cholesterol, mg/dL | 215.6 (44.8) | 214.3 (40.4) | 197.6 (35.7) | 191.3 (36.5) | 199.4 (38.2) | 193.0 (31.5) | 224.6 (43.9) | 234.0 (39.4) | 226.6 (41.1) | 200.1 (43.5) |
| Systolic blood pressure, mmHg | 128.2 (20.9) | 118.0 (16.8) | 121.6 (19.8) | 129.6 (21.8) | 123.9 (21.0) | 121.5 (20.3) | 130.6 (17.0) | 128.9 (17.4) | 126.4 (17.5) | 121.5 (17.7) |
| Diastolic blood pressure, mmHg | 79.8 (12.0) | 71.6 (10.0) | 70.1 (10.0) | 74.2 (10.1) | 71.0 (10.1) | 71.3 (10.3) | 78.5 (8.9) | 75.1 (9.0) | 75.5 (9.2) | 73.1 (10.8) |

AA: African American, CHN: Chinese-Ancestry, EA: European-American, HIS: Hispanic/Latinos

ARIC: Atherosclerosis Risk in Communities Study, HCHS/SOL: Hispanic Community Health Study/Study of Latinos, MESA: Multi-Ethnic Study of Atherosclerosis, PAGE: Population Architecture using Genetic Epidemiology study, WHI: Women’s Health Initiative.

| **Table S5**. **Partial correlations between six contiguous traits decomposing the ECG**, in n=10,618 eligible Hispanic Community Health Study/Study of Latinos participants. Partial correlations were adjusted for RR interval, gender, study site, and ancestry principal components. | | | | | | |
| --- | --- | --- | --- | --- | --- | --- |
|  | P wave | PR segment | QRS interval | ST segment | T wave | TP segment |
| P wave | 1 | -0.27 | 0.17 | -0.02 | 0.01 | -0.59 |
| PR segment | -0.27 | 1 | -0.04 | -0.04 | -0.07 | -0.37 |
| QRS interval | 0.17 | -0.04 | 1 | -0.04 | -0.27 | -0.20 |
| ST segment | -0.02 | -0.04 | -0.04 | 1 | -0.20 | 0.00 |
| T wave | 0.01 | -0.07 | -0.27 | -0.20 | 1 | -0.47 |
| TP segment | -0.59 | -0.37 | -0.20 | 0.00 | -0.47 | 1 |

| **Table S6.** **Published genome-wide association studies of electrocardiographic traits**, indexed on the NHGRI GWAS catalog (Sept. 30, 2018). | | | | | |
| --- | --- | --- | --- | --- | --- |
| **Date** | **PMID** | **First author** | **# new loci** | **Study title** | **N (discovery)** |
| **P wave** |  |  |  |  |  |
| 8/1/2017 | 28794112 | Christophersen IE | 20 | Fifteen Genetic Loci Associated With the Electrocardiographic P Wave. | EUR: 3,7678  AFR: 6,778 |
| 5/21/2014 | 24850809 | Verweij N | 3 | Genetic determinants of P wave duration and PRseg segment. | EUR: 16,468 |
| **PR interval** |  |  |  |  |  |
| 1/10/2010 | 20062060 | Pfeufer A | 9 | Genome-wide association study of PRseg interval. | 28,517 |
| 1/10/2010 | 20062063 | Holm H | 4 | Several common variants modulate heart rate, PRseg interval and QRS duration. | 10,000 |
| 2/10/2011 | 21347284 | Smith JG | 5 | Genome-wide association studies of the PRseg interval in African Americans. | AFR:  6,247 |
| 11/8/2012 | 23139255 | Butler AM | 8 | Novel loci associated with PRseg interval in a genome-wide association study of 10 African American cohorts. | AFR: 13,415 |
| 11/10/2017 | 29127183 | Seyerle AA | 8 | Genome-wide association study of PRseg interval in Hispanics/Latinos identifies novel locus at ID2 | H/L: 14,756 |
| 7/17/2014 | 25035420 | Hong KW | 3 | Identification of three novel genetic variations associated with electrocardiographic traits (QRS duration and PRseg interval) in East Asians. | EAS: 6805 |
| 1/10/2010 | 20062061 | Chambers JC | 1 | Genetic variation in SCN10A influences cardiac conduction. | EAS: 6,243  EUR: 5,370 |
| In press | * | Wojcik, G | 5 | The PAGE Study: How Genetic Diversity Improves Our Understanding of the Architecture of Complex Traits | AFR: 3,360  H/L: 13,702 |
| 7/25/2018 | 30046033 | Van Setten, J | 50 | PR interval genome-wide association meta-analysis identifies 50 loci associated with atrial and atrioventricular electrical activity. | EUR: 92,000 |
| **PR segment** |  |  |  |  |  |
| 5/21/2014 | 24850809 | Verweij N | 8 | Genetic determinants of P wave duration and PRseg segment. | EUR: 16,468 |
| **QRS interval** |  |  |  |  |  |
| 7/17/2014 | 25035420 | Hong KW | 3 | Identification of three novel genetic variations associated with electrocardiographic traits (QRS duration and PRseg interval) in East Asians. | EAS: 6805 |
| 9/27/2016 | 27659466 | van der Harst P | 44 | 52 Genetic Loci Influencing Myocardial Mass. | EUR: 73,518 |
| 8/29/2016 | 27577874 | Evans DS | 42 | Fine-mapping, Novel Loci Identification, and SNP Association Transferability in a Genome-Wide Association Study of QRS Duration in African Americans. | AFR: 13,031 |
| 11/14/2010 | 21076409 | Sotoodehnia N | 12 | Common variants in 22 loci are associated with QRS duration and cardiac ventricular conduction. | EUR: 40,407 |
| 1/10/2010 | 20062063 | Holm H | 3 | Several common variants modulate heart rate, PRseg interval and QRS duration. | EUR: 10,000 |
| 3/5/2013 | 23463857 | Ritchie MD | 5 | Genome- and phenome-wide analyses of cardiac conduction identifies markers of arrhythmia risk. | EUR: 5,272 |
| In press | * | Wojcik, G | 5 | The PAGE Study: How Genetic Diversity Improves Our Understanding of the Architecture of Complex Traits | AFR: 3,274  H/L: 13,420 |
| 12/13/2016 | 27958378 | Floyd JS | 1 | Large-scale pharmacogenomic study of sulfonylureas and the QT, JT and QRS intervals: CHARGE Pharmacogenomics Working Group. | EUR: 45 002  AFR: 11,731  H/L: 15,124 |
| **QT interval** |  |  |  |  |  |
| 7/23/2014 | 25055868 | Sano M | 1 | Genome-wide association study of electrocardiographic parameters identifies a new association for PR interval and confirms previously reported associations. | EAS: 2994 |
| 6/22/2014 | 24952745 | Arking DE | 68 | Genetic association study of QT interval highlights role for calcium signaling pathways in myocardial repolarization. | EUR: 76,061  AFR: 13,105 |
| 3/22/2009 | 19305409 | Pfeufer A | 12 | Common variants at ten loci modulate the QT interval duration in the QTSCD Study. | EUR: 15,842 |
| 3/22/2009 | 19305408 | Newton-Cheh C | 13 | Common variants at ten loci influence QT interval duration in the QTGEN Study. | EUR: 13,685 |
| 8/1/2009 | 20031603 | Marroni F | 2 | A genome-wide association scan of RR and QT interval duration in 3 European genetically isolated populations: the EUROSPAN project. | EUR: 2325 |
| 11/19/2012 | 23166209 | Smith JG | 2 | Impact of ancestry and common genetic variants on QT interval in African Americans. | AFR: 13,105 |
| 7/9/2009 | 19587794 | Nolte IM | 2 | Common genetic variation near the phospholamban gene is associated with cardiac repolarisation: meta-analysis of three genome-wide association studies. | EUR: 29,400 |
| 1/10/2010 | 20062063 | Holm H | 4 | Several common variants modulate heart rate, PRseg interval and QRS duration. | EUR: 10,000 |
| 4/30/2006 | 16648850 | Arking DE | 1 | A common genetic variant in the NOS1 regulator *NOS1AP* modulates cardiac repolarization. | EUR: 3,966 |
| 6/20/2012 | 22726844 | Kim JW | 2 | A common variant in *SLC8A1* is associated with the duration of the electrocardiographic QT interval. | EAS: 6,805 |
| In press | * | Wojcik, G | 5 | The PAGE Study: How Genetic Diversity Improves Our Understanding of the Architecture of Complex Traits | AFR: 3,293  H/L: 13,698 |
| 12/13/2016 | 27958378 | Floyd JS | 2 | Large-scale pharmacogenomic study of sulfonylureas and the QT, JT and QRS intervals: CHARGE Pharmacogenomics Working Group. | EUR: 45 002  AFR: 11,731  H/L: 15,124 |
| 1/10/2010 | 20062061 | Chambers JC | 1 | Genetic variation in *SCN10A* influences cardiac conduction. | EAS: 6,543 |

*Article in press in Nature. Preprint version available on bioRxiv: <https://www.biorxiv.org/content/10.1101/188094v2>

**Table S7**. **Previously-reported associations of SNPs with temporal electrocardiographic traits**, and corresponding results in N=34,668 participants in the Population Architecture using Genomics and Epidemiology (PAGE) study and the Multiethnic Study of Atherosclerosis (MESA).

Table S7 is provided in a separate excel spreadsheet.

| **Table S8 Trait-specific direction of effects, and meta-analysis heterogeneity p-values for univariate associations (p_univariate_ < 5×10^-9^)** **of loci discovered in**  **combined-phenotype analyses** **of six electrocardiographic traits (p_aSPU_ < 5x10^-9^)**, in 34,668 participations from the Population Architecture using  Genomics and Epidemiology study (PAGE) and Multi-Ethnic Study of Atherosclerosis (MESA). | | | | | | | | | | | | | | | |
| --- | --- | --- | --- | --- | --- | --- | --- | --- | --- | --- | --- | --- | --- | --- | --- |
|  |  |  |  | P wave | | PR Segment | | QRS Interval | | ST Segment | | T wave | | TP Segment | |
| SNP | Chr | Position | Locus | Dir | Het p | Dir | Het p | Dir | Het p | Dir | Het p | Dir | Het p | Dir | Het p |
| rs13143308 | 4 | 111714419 | PITX2 | +++ | 0.004 | +-- | 0.447 |  |  | +++ | 0.945 |  |  |  |  |
| rs4340921 | 5 | 49687697 | EMB | --- | 0.557 | ++- | 0.916 |  |  |  |  |  |  | -++ | 0.518 |
| rs3211938 | 7 | 80300449 | CD36 | -??- | 0.661 | -??- | 0.706 | -??+ | 0.03 |  |  | -??- | 0.009 | +??+ | 0.124 |
| rs11073663 | 15 | 85260268 | ZNF592 |  |  | +-- | 4×10^-4^ |  |  |  |  | +++ | 0.527 | -++ | 0.016 |
| rs142166837 | 17 | 57471022 | YPEL2 |  |  |  |  | --- | 0.834 |  |  | +++ | 0.509 | --- | 0.331 |
| rs13047360 | 21 | 28851580 | BC043580 |  |  | --- | 0.340 | +++ | 0.017 |  |  |  |  |  |  |

Directions (+ or -) listed for ancestry groups, in the order: African Americans, European-American, Chinese Ancestry, Hispanic/Latinos.

Question marks (?) indicate that SNPs were missing, or filtered out for the corresponding ancestry group

Cells are left blank if the association p-value is not nominally significant (p<0.05).

| **Table S9 Trait-specific trans-ethnic meta-analyzed effect estimates and standard errors for univariate associations** **(p_univariate_ < 5×10^-9^) of loci**  **discovered in combined-phenotype analyses of six electrocardiographic traits (p_aSPU_ < 5x10^-9^)**, in 34,668 participations from the Population Architecture  using Genomics and Epidemiology study (PAGE) and the Multi-Ethnic Study of Atherosclerosis (MESA). | | | | | | | | | | | |
| --- | --- | --- | --- | --- | --- | --- | --- | --- | --- | --- | --- |
|  |  |  |  | Effect size (standard error) | | | | | | | |
| SNP | Chr | Position | Locus | P wave | PR Segment | QRS Interval | ST Segment | T wave | TP Segment | QT interval | PR interval |
| rs13143308 | 4 | 111714419 | PITX2 | **0.6 (0.09)** | -0.6 (0.17) | 0.1 (0.07) | 0.1 (0.07) | -0.2 (0.15) | -0.4 (0.24) | 0.1 (0.19) | 0 (0.15) |
| rs4340921 | 5 | 49687697 | EMB | **-0.6 (0.08)** | 0 (0.15) | 0.2 (0.06) | 0 (0.06) | -0.2 (0.13) | 0.8 (0.21) | 0.1 (0.17) | -0.5 (0.13) |
| rs3211938 | 7 | 80300449 | CD36 | -1.4 (0.34) | -1.7 (0.65) | -0.7 (0.26) | 0.3 (0.29) | -2.6 (0.58) | **6.7 (0.9)** | **-3.7 (0.74)** | -3.4 (0.6) |
| rs11073663 | 15 | 85260268 | ZNF592 | 0 (0.09) | **1 (0.16)** | -0.1 (0.07) | 0 (0.07) | 0.2 (0.14) | -1.2 (0.23) | 0.2 (0.18) | 1 (0.14) |
| rs142166837 | 17 | 57471022 | YPEL2 | 0.1 (0.08) | -1 (0.15) | 0 (0.06) | -0.1 (0.06) | **0.4 (0.13)** | 0.6 (0.21) | 0.3 (0.17) | -0.8 (0.13) |
| rs13047360 | 21 | 28851580 | BC043580 | -0.2 (0.08) | 0.1 (0.15) | **-0.2 (0.06)** | 0.1 (0.06) | 0.9 (0.13) | -0.5 (0.21) | 0.7 (0.17) | 0 (0.13) |

Bolded values are significant at the genome-wide level (p_univariate_<5×10^-9^)

**Table S10 Coded allele frequencies and imputation quality of lead variants for loci discovered in combined-phenotype analyses of six electrocardiographic traits (p_aSPU_ < 5x10^-9^)**, in 34,668 participations from the Population Architecture using Genomics and Epidemiology study (PAGE) and the Multi-Ethnic Study of Atherosclerosis (MESA).

|  | **Coded Allele Frequency** | | | | **ARIC** | | **WHI (EA)** | | | | | **PAGE** |
| --- | --- | --- | --- | --- | --- | --- | --- | --- | --- | --- | --- | --- |
| SNP | AA | CHN | EA | HIS | AA | EA | GARNET | GECCO | HIPFX | MOPMAP | WHIMS |  |
| rs4340921 | 66% | 46% | 49% | 44% | 0.995 | 0.995 | 0.995 | 0.983 | 0.988 | 0.987 | 0.995 | 0.978 |
| rs13143308 | 30% | 21% | 26% | 39% | 0.987 | 0.997 | 0.999 | 0.995 | 0.998 | 0.999 | 0.998 | 1 |
| rs11073663 | 27% | 54% | 19% | 48% | 0.988 | 0.996 | 1 | 0.998 | 0.998 | 0.993 | 1 | 1 |
| rs142166837 | 31% | 52% | 32% | 49% | 0.993 | 0.988 | 0.99 | 0.99 | 0.991 | 0.99 | 0.992 | 1 |
| rs3211938 | 10% | 1% | <0.01% | <0.01% | 0.896 | 0.886 | * | * | * | * | * | 0.992 |
| rs142166837 | 19% | 38% | 32% | 45% | 0.937 | 0.979 | 1 | 0.922 | 1 | 0.835 | 0.933 | 0.994 |

AA: African American, CHN: Chinese-American, EA: European-American, HIS: Hispanic/Latinos

ARIC: Atherosclerosis Risk in Communities Study, HCHS/SOL: Hispanic Community Health Study/Study of Latinos, MESA: Multi-Ethnic Study of Atherosclerosis, PAGE: Population Architecture using Genetic Epidemiology study, WHI: Women’s Health Initiative.

PAGE analysis sample includes African-American and Hispanic participants from the Women’s Health Initiative, and all participants genotyped in the Hispanic Community Study/Study of Latinos.

* Major allele frequency below filtering threshold (<1%)
